# Differences in growth features between species are driving cereal-legume intercrop yield: a statistical learning approach based on aggregated dataset

**DOI:** 10.1101/2024.09.17.613461

**Authors:** Rémi Mahmoud, Noémie Gaudio, Xavier Gendre, Nadine Hilgert, Pierre Casadebaig

## Abstract

Increasing crop diversification is crucial for developing more sustainable agricultural systems, and cereal-legume intercropping is a promising strategy. This study investigates the factors influencing the yield of cereal-legume intercrops using data from six field experiments in southwestern France, where durum wheat was intercropped with either faba bean or pea. We assessed how differences in plant traits between the associated species (*e.g.*, height or biomass growth rates) are related to the intercrop productivity. Additionally, we developed a novel modeling approach, combining machine learning and mixed-effects models, to identify the key traits driving intercrop performance based on variable importance.

Our results show that interspecific differences in plant traits, particularly in biomass accumulation rate, maximum leaf area index, and elongation rate, were the most important factors explaining intercrop yield. These traits and their differences mainly suggest that competitive processes shape the outcome of a mixture and highlight the importance of dynamic measurements in agronomic experiments. The relationship between species yield and trait differences was symmetric for both intercropped species. Furthermore, these relationships were scale-dependent, with trends observed at the aggregate level not always consistent at the level of individual experiments.

Our study highlights the importance of considering trade-offs when designing intercropping systems for practical applications and demonstrates the value of combining machine learning with ecological knowledge to gain insights into complex agricultural systems from aggregated datasets.

**Highlights:**

- Joint analysis of experimental datasets provided new insights into crop mixtures functioning.
- Differences in growth traits between species in the mixture predicted their performance.
- The strength of the correlation between performance and trait distance was similar for both species.
- Farmers and researchers need to consider trade-offs when designing intercropping systems.

## Introduction

Increasing crop diversification is a key practice towards more sustainable agricultural systems (Duru et al., 2015), with overall beneficial effects observed (Beillouin et al., 2021). At the annual and field scale, an example of a well-studied diversification strategy is inter-cropping, *i.e.*, combining two different species in the same field for most of their growing periods. Since different species differ in their ecophysiological functioning, growing a mixture could improve resource use efficiency relative to the species grown separately in sole crops. Particularly, cereal-legume mixtures are a highly effective practice, not only because of the complementary nitrogen use of the two species (Landschoot et al., 2024), but also because of their practical feasibility (Verret et al., 2020). Meta-analyses generally indicate that intercropping increases productivity per unit area in low-input contexts (Bedoussac et al., 2015). However, results from field experiments suggest that these benefits are highly context-dependent (Jones et al., 2023), with some situations showing increased productivity and others showing decreased productivity compared to sole cropping (Martin-Guay et al., 2018; MacLaren et al., 2023).

This discrepancy between broad frameworks linking diversity to productivity (Brooker et al., 2021) and local outcomes calls for a focus on the mechanisms behind the effects of diversity, specifically whether it acts as insurance (Loreau et al., 2021) or through better resource use complementarity between species. The underlying idea is that it is not diversification *per se* that leads to productivity gains but rather the choice of species to combine in a given environment to promote positive plant-plant interactions while minimizing negative ones (Dee et al., 2023; McGuire, 2023). In this context, understanding how species characteristics impact mixture productivity is crucial for developing intercropping as a component of more sustainable systems (Wang et al., 2024b).

Modeling is a relevant tool for improving our understanding of systems, especially for assessing how species characteristics are related to mixture productivity. As pointed out by van Ooyen (2011), *most formal models lie on a continuum between two extreme categories: mechanistic models and phenomenological models*. Mechanistic models are based on in-depth knowledge of ecophysiological mechanisms and include numerous parameters with explicit biological meanings. In contrast, phenomenological models start from experimental data to explore correlations arising from potential causal relationships. They do not require *a priori* knowledge of the biological mechanisms underlying the observations, but rely heavily on data availability (Gunawardena, 2014). Within the phenomenological models, Breiman (2001a) highlighted a distinction between classical statistical models (such as generalized linear models) and algorithmic models (*e.g.*, random forests and neural networks). Algorithmic models are valued for their flexibility in learning patterns from data without relying on assumptions about specific data distributions, making them particularly well-suited for handling complex biological data. Additionally, their ability to assess the contribution of different inputs to the studied process provides a distinct advantage in the analysis of biological systems.

When it comes to modeling species mixtures, mechanistic models are often limited in the number of crop species they can represent and in their ability to capture plant-plant interactions and phenotypic plasticity (Gaudio et al., 2019), while these processes are critical to understanding mixture performance (Mahaut et al., 2023). As a result, we opted for an algorithmic modeling approach in which plant-plant interactions are inferred from observed data rather than explicitly described from an ecophysiological perspective. In this context, differences between species characteristics, commonly referred to as plant traits (Volaire et al., 2020), are valuable for elucidating plant-plant interactions (Kunstler et al., 2012) and are good proxies for estimating both the overall performance of the mixture and the performance of each species within it (Montazeaud et al., 2018; Gaudio et al., 2021; Mahaut et al., 2023). Depending on the trait considered, species differences may reflect either competition or complementarity (Wagg et al., 2017). For example, competition has been associated with differences in maximum height (Carmona et al., 2019) or early growth rates (Zhang et al., 2020), while niche complementarity has been associated with differences in root length (Homulle et al., 2021). Although species differences are expected to influence mixture functioning positively (through facilitation and/or complementarity) or negatively (through competition), the strength and shape of this relationship in agricultural systems remain largely unknown (Gaudio et al., 2021; Mahaut et al., 2023). We argue that using ecological concepts and agronomic knowledge to approximate plant-plant interactions, rather than working directly from the raw observed phenotypes, would increase the interpretability of our model.

In this study, we therefore approximated plant-plant interactions as a function of species trait differences to predict the yield of each species in the mixture and to assess the relative contribution of each trait to species yield. We used the results of six field experiments conducted in southwestern France that included species-dependent measurements along with mixture and sole crop productivity. The experiments included two wheat/legume intercrops, *i.e.*, durum wheat (*Triticum turgidum* L.) associated with faba bean (*Vicia faba* L.) or pea (*Pisum sativum* L.). We summarized plant-plant interactions using relative trait distance (Kunstler et al., 2012) under two conditions: (1) within the mixture, between species, *i.e.*, interspecific indicators (Gaudio et al., 2021), and (2) within a species, between sole-and intercropping conditions, *i.e.*, intraspecific indicators (Engbersen et al., 2022). The choice of relative distance over absolute distance was primarily determined by the nature of the traits measured in agronomic experiments (height, biomass, leaf area index), which mainly reflect asymmetric plant-plant interactions, and thus competition (Mahaut et al., 2023). Also called fitness differences, they are assumed to characterize the competitive advantage of one species over another one, for example due to asymmetry in species growth rates (Wang et al., 2024b).

We then developed a modeling approach that combines machine learning and inference, as used in previous studies in ecology (Yang et al., 2022; Primka IV et al., 2023), based on the random forest algorithm and linear mixed-effects model. We also compared this original combined approach with each of the individual methods involved. To interpret the model outputs and identify important features for mixture performance, we used a two-step process. First, we reduced the set of features through a rigorous variable selection during model fitting. Then, we ranked the selected variables by estimating their contribution to the variance in mixture performance (variable importance).

**Figure 1.**
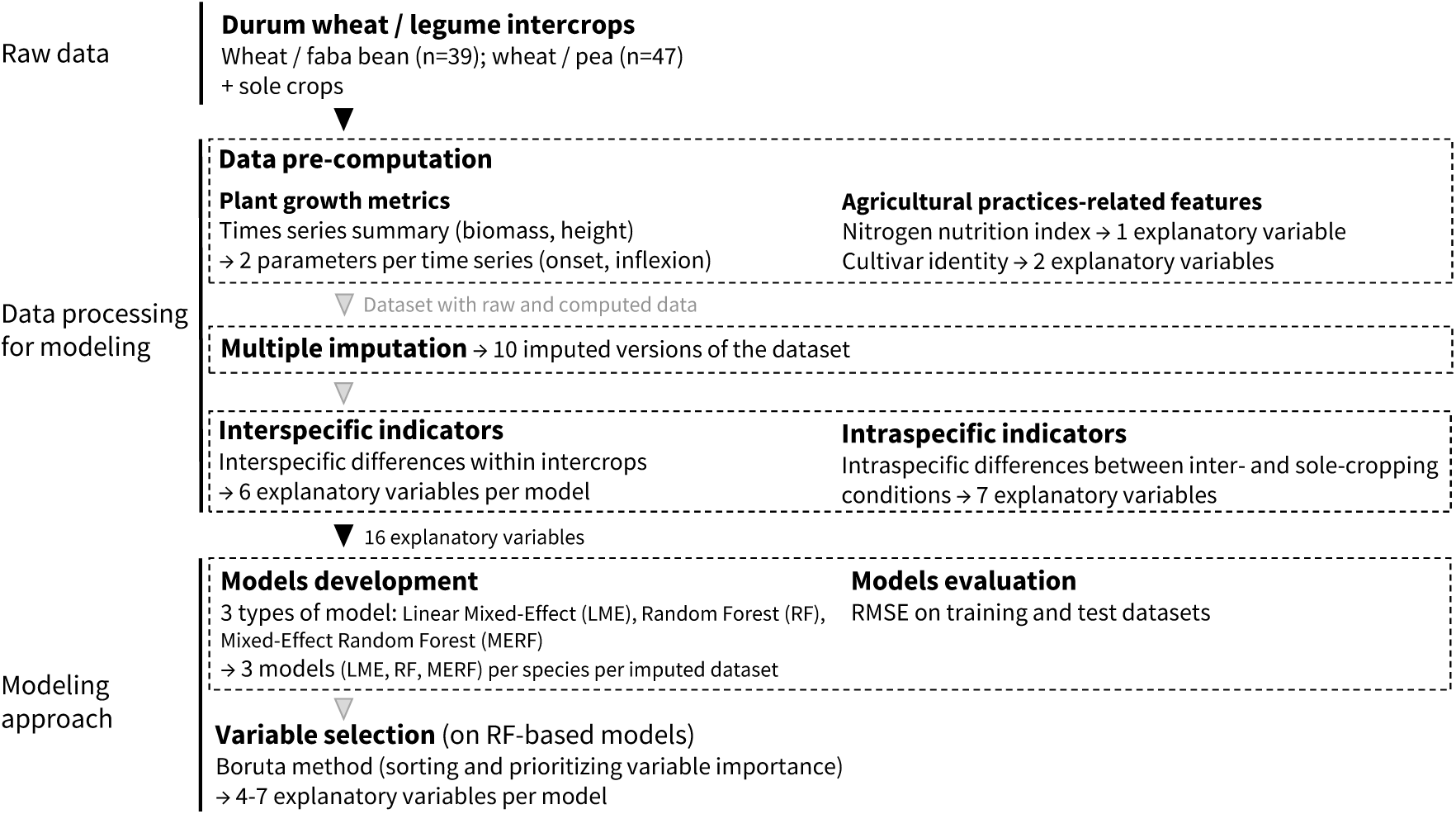
Description of the main steps of the modeling workflow. Starting with a dataset including two mixtures (*i.e.*, durum wheat associated with faba bean or pea), we first summarized the plant growth dynamics and the effect of agricultural practices. We then applied an imputation procedure to obtain 10 imputed versions of the dataset. We summarized plant-plant interactions under two conditions: (1) within the mixture, between species (*i.e.*, interspecific indicators), and (2) within a species, between sole-and intercropping conditions (*i.e.*, intraspecific indicators). Finally, we fitted the yield of each species in a mixture to these features using a random forest model.

## Data processing for modeling

### Raw data: Field experiments and plant measurements

The raw dataset gathers the results of 6 field experiments involving annual and synchronous (*i.e.*, crop species sown and harvested at the same time) cereal-legume intercrops and their sole crop reference. The experiments were selected from an open-source global dataset (Gaudio et al., 2023) as a compromise between the number of common variables across experiments and the number of intercrops we could consider. For an experiment to be selected, we required that growth-related variables were measured dynamically during the crop cycle and that it included common species across experiments.

The field experiments were carried out in southwestern France, covering 6 years from 2006 to 2013. The year 2013 stands out from the others because of the high amount of precipitation recorded during the growing season (Table 1). The dataset includes one cereal species, *i.e.*, durum wheat (*Triticum turgidum* L.); two legume species, *i.e.*, faba bean (*Vicia faba* L.) and pea (*Pisum sativum* L.); and the two resulting winter wheat/legume intercrops. The experiments also differed in terms of the cultivars used. For wheat/faba bean mixtures, 2010 and 2011 were almost similar, with the same wheat cultivar and different faba bean cultivars each year (Table 1). For wheat/pea mixtures, 2006 and 2007 included the same cultivars for both pea and wheat (Bedoussac and Justes, 2010), with 3 additional wheat cultivars included in 2007. The years 2012 and 2013 were more diverse, with 3 wheat cultivars tested each year, along with 4 faba bean cultivars and 4-5 pea cultivars that differed in height and earliness (Kammoun et al., 2021).

**Table 1.** Main characteristics of the data used, coming from 6 experiments conducted in southwestern France. Two cereal/legume intercrops were involved, *i.e.*, wheat/faba bean and wheat/pea, with 1-5 cultivars per species and per experimental year, with the two intercrops being studied within the same experiment in 2012 and 2013. The environment was characterized by the mean temperature (°C) and the sum of precipitation (mm) recorded from crop sowing to harvest. Within each experiment, several experimental units (unique combination of {year, crop management}) were involved for each mixture.

| Year | Intercrop | Precipitations<br>(mm) | Temperature<br>($^{\circ}\text{C}$) | n | No. cultivar<br>wheat | No. cultivar<br>legume |
| --- | --- | --- | --- | --- | --- | --- |
| 2006 | wheat / pea | 473.2 | 10.0 | 3 | 1 | 1 |
| 2007 | wheat / pea | 546.4 | 11.4 | 16 | 4 | 1 |
| 2010 | wheat /<br>fababean | 501.4 | 10.1 | 16 | 1 | 1 |
| 2011 | wheat /<br>fababean | 311.9 | 10.7 | 4 | 1 | 1 |
| 2012 | wheat /<br>fababean | 435.6 | 10.2 | 9 | 3 | 4 |
| 2012 | wheat / pea | 435.6 | 10.2 | 9 | 3 | 4 |
| 2013 | wheat /<br>fababean | 782.6 | 10.4 | 10 | 3 | 4 |
| 2013 | wheat / pea | 782.6 | 10.4 | 19 | 3 | 5 |

In total, the dataset contains 86 experimental units of species mixtures, defined as the unique combination of {year, management} levels, with the crop management including species and cultivar choice as well as agricultural interventions (especially different levels of N-fertilization). Among these 86 experimental units, 39 and 47 were represented by wheat/faba bean and wheat/pea intercrops, respectively (Table 1). Within the six experiments, several plant variables were measured at the experimental unit scale. Grain yield (t.ha^-1^) was measured systematically. Crop height (m) and aboveground biomass (t.ha^-1^) were measured dynamically throughout the growing season. Leaf area index (LAI, leaf area per soil area, in m^2^.m^-2^) and specific leaf area (SLA, cm^2^.g^-1^) were measured at their maximum value, and nitrogen (N) content in aboveground biomass (%) at flowering, allowing the calculation of the N nutrition index (NNI, Louarn et al., 2021). Given the heterogeneity of the gathered experiments, we used a multiple data imputation procedure to handle the small fraction of missing data.

### Data pre-computation: plant growth metrics and nitrogen status

#### Plant growth metrics

Shoot biomass and crop height were measured dynamically during the cropping season in each experimental unit, with measurement dates varying between experiments. Assuming that plant growth followed a logistic curve, we summarized the process with three common parameters having a biological meaning (Zwietering et al., 1990). We used smoothing splines to reduce these curves’ dimensions by extracting two parameters: i) the slope at the inflexion point (*i.e.*, growth rate) and ii) the onset of the growth phase. Given the coordinates (*x, y*) at the inflexion point, the onset was estimated as the intersection of the tangent at the inflexion point with the x-axis. We did not estimate these parameters if the measurements were not done over the entire growth cycle. Additionally, the maximum values were also directly extracted for three plant variables: height, LAI and SLA. We chose to exclude maximum biomass from the further calculation of explanatory variables due to its high positive correlation with yield, as illustrated in studies focusing on reproductive allometry (Gaudio et al., 2021).

#### Nitrogen status of intercropped species

To assess individual species N status in mixture, we calculated the Nitrogen Nutrition Index (NNI; Louarn et al. (2021)). The critical N dilution curve is species-specific and describes the minimum aboveground N concentration required to achieve maximal growth for a given biomass (*Nc*, in %; Justes et al. (1994)). The NNI evaluates a crop N status by comparing its N concentration to *Nc*. NNI above (below) 1 indicates N excess (stress) in the crop (Lemaire and Meynard, 1997). NNI was computed at flowering for each species as following (Louarn et al., 2021):

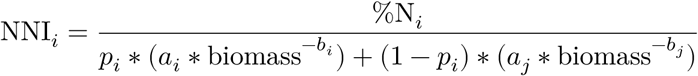

where *a_i_*, (*a_j_*) and *b_i_* (*b_j_*) are parameters for species *i* (resp. *j*), %N*_i_* is the N concentration of species *i* and *p_i_* is the proportion of species within the intercrop. The parameters *a* (%) and *b* (unitless) are specified identically for pea and faba bean with *a* at 5.1 % and *b* at 0.32 (Louarn et al., 2021), as well as for wheat with *a* at 5.4 % and *b* at 0.44 (Justes et al., 1994).

### Data imputation

Given that the six gathered experiments did not follow strictly identical protocols, some data were missing, resulting in an unbalanced dataset (Mahmoud et al., 2024). Listwise data deletion is a common procedure that retains only observations with complete entries in a dataset, but it may drastically reduce the sample size and bias the analysis (Roth, 1994). Alternatively, the multiple imputation allows getting multiple plausible values for each missing entry, while keeping track of the variability due to the imputation process (Schafer, 1999).

To perform multiple imputation, we used the JointAI R package algorithm (Erler et al., 2021), which is based on a Bayesian mixed-effects model that allowed tracking the within-experiment dependence between observations. This method provided 10 imputed versions of the dataset.

### Explanatory variables selected and computed for modeling

We aimed to model the yield of each species in each mixture (wheat/faba bean and wheat/pea), resulting in four distinct models. Each model is based on a common set of 16 explanatory variables, related to plant characteristics, their interactions, and the effect of agricultural practices.

#### Plant-related explanatory variables

Guided by ideas from community ecology, we computed two sets of plant-related indicators at the experimental unit scale (Table 2) from the data processed in the previous steps. Within each intercrop, the difference in a given plant variable between the cereal and the legume was calculated as a proxy for interspecific plant-plant interactions (*i.e.*, interspecific indicators), resulting in 6 shared explanatory variables per species. Within each crop species, the difference in a given plant variable between intercrop and sole crop indicated the potential effect of the mixture context (*i.e.*, intraspecific indicators). This resulted in 7 explanatory variables per species, which differed between wheat and legumes. As these differences are symmetric depending on the focus species, only legume intraspecific differences were included in legumes models, and similarly for wheat models. Regarding maximum SLA, we opted to exclude its calculation from explanatory variables for interspecific differences, while retaining it for intraspecific ones. This plant trait did not seem relevant to compare cereals and legumes given their highly contrasting leaf morphology. Then, SLA intraspecific variability makes more sense than interspecific difference (Lisner et al., 2021) in terms of interpretability.

**Table 2.** List of the plant-related explanatory variables used as input in the models in order to explain yield of intercropped species. For the different variables (biomass, height, LAI, and SLA), 1-3 parameters were available to characterize plant growth and stature. Two sets of indicators were computed: i) interspecific indicators were calculated within each intercrop as the difference for a plant parameter between the cereal and the legume; ii) intraspecific indicators were calculated for a given species as the difference for each plant parameter between intercrop and sole crop.

| Plant Variables | Parameter | Unit | Interspecific<br>($x_{cereal} - x_{legume}$) | Intraspecific<br>($x_{intercrop} - x_{sole\ crop}$) |
| --- | --- | --- | --- | --- |
| Biomass | Onset | $^{\circ}C.d$ | X | X |
| | Inflexion | $t.ha^{-1}$ | X | X |
| Height | Onset | $^{\circ}C.d$ | X | X |
| | Inflexion | $m.^{\circ}Cd-1$ | X | X |
| | Max | $m$ | X | X |
| Leaf area index (LAI) | Max | $m^2.m^{-2}$ | X | X |
| Specific leaf area (SLA) | Max | $cm^2.g^{-1}$ | - | X |

#### Agricultural practices-related explanatory variables

To summarize agricultural practices, we used two sets of indicators. The first one simply corresponds to the cultivar identity. The cultivar identity of both intercropped species was included in each species model (Table 1), resulting in two additional explanatory variables per model. The second agronomic indicator was the crop N status (NNI). In each species model, only the NNI of the considered species was included as an explanatory variable.

## Modeling approach

### Objectives and strategy

We start the modeling step from the imputed datasets presented in the previous section, each with *p* = 16 explanatory variables. To study how the environment affected mixture functioning, we considered *experiments* as a random effect (and therefore not count this factor as an explanatory variable, rather as an indicator vector).

Our main issue was to identify, among these variables, which ones best explain the yield of species in mixture, and which modeling approach is most relevant to understand these systems. The choice of the modeling strategy then depends largely on these data and their dependency structure. Here, the data came from *l* = 6 experiments, with 4 experiments per intercrop, containing *n* = 39 and *n* = 47 observations per mixture (wheat/faba bean and wheat/pea, respectively; Table 1). The fact that the observations were partly drawn from common experiments resulted in a distinct grouped structure and dependency within the observations, thereby challenging the classical independence assumption of many statistical learning algorithms (Hastie et al., 2009).

We set up and compared three modeling approaches: i) random forest (RF) algorithm ii) linear mixed-effects (LME) model, and iii) mixed-effects random forest (MERF) algorithm, which combines both previous approaches (Hajjem et al., 2012; Capitaine et al., 2021). We mainly focused on the MERF approach, as we assume that this approach takes advantages of both machine learning and mixed-effects models, *i.e.* handles both the unknown and potentially non-linear relationships between predictors and outputs, and the ability to interpret the *experiments* effect within the residual error (within-group observation dependencies).

The RF algorithm allows the aggregation of predictions obtained from a large number of regression trees trained on different subsets of variables (Breiman, 2001b). It provides high prediction quality and can capture interactions between explanatory variables. This method is here proposed as baseline, as it is a widely used approach and known to be suitable for a large range of problems.

Statistical LME models are an extension of the common linear models to include both fixed and random effects, taking into account the nested structure of the data and the within-group dependency of the observations. More than a simple baseline, the comparison between the LME and the MERF models will allow us to test our assumption that accounting for non-linearity in predictions matters.

Although we reported about the performance of the three modeling approaches, we will focus on the agroecological significance of results from the MERF algorithm. In the following section, we choose to write models as loss functions, *i.e.* the sum of the squared differences between the actual and predicted values. We used this form as a common way to quantify the error of a prediction model, whether it is algorithmic or statistical.

The LME approach is well-known and widely used. It could be used to explain the yield *y* ϵ *ℝ* as the sum of fixed effects given by a linear combination *β*^T^*x* of the *p* explanatory variables *x* = (*x*^1^,…,*x^p^*)^T^ ϵ ℝ*^p^* and random effects *u*^T^*z* where *z* = (*z*^1^,…,*z^l^*)^T^ ϵ {0,1}*^l^* stands for the experiment indicator vector that gives the group membership of the experiment. To fit the model parameters *β* ϵ ℝ*^p^* and *u* ∈ ℝ*^l^*, the following least squares criterion is minimized,

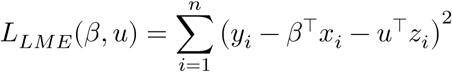

where *y*_*i*_ is the yield observed in the experimental unit *i* ϵ {1, …, *n*} with explanatory variables *x*_*i*_ and indicator variables *x*_*i*_.

The relationship between the yield and the variables is assumed to be linear in the LME approach. Such an hypothesis is debatable in practice and non-linearity is often relevant and desirable. To this end, a non-parametric (with no assumption on the relationship shape) model such as RF is a commonly used alternative. RF explains *y* with an aggregation of decision trees 𝒯 represented hereafter by a function *g*_𝒯_. Models were fitted with the randomForest R package (Liaw and Wiener, 2002). To take into account both explanatory and indicator variables, the least squares criterion has to be minimized with respect to the decision tree collection 𝒯,

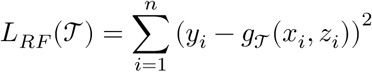

The drawback of the pure RF approach is that the explanatory variables and experiment indicator are both integrated in the model. It is no longer feasible to identify the contribution of *z* as a random effect and to interpret the result in this sense. To circumvent this and retrieve random effects, the MERF approach consists of mimicking the form of LME but replacing the fixed effects term with an RF model based on the explanatory variables only. MERF models were fitted using the LongituRF R package (Capitaine, 2020). The least squares criterion to be minimized is then

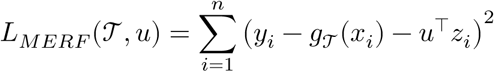

### Variable selection in random forest-based models

Basically, the RF algorithm does not include a variable selection procedure, whereas it would allow for greater parsimony, by explaining the response variable with a limited set of explanatory variables (by discarding non-informative variables), eliminating redundant variables, and facilitating interpretation. The issue of parsimony is significant in our case, as we introduced 16 explanatory variables in each model. Therefore, the RF algorithm was extended with a variable selection procedure, based on the Boruta method within the Boruta R package (Kursa and Rudnicki, 2010a), which has been shown to be robust and interpretable (Speiser et al., 2019). The Boruta method consists in creating shuffled duplicates of each explanatory variable. The importance of each variable (raw and shuffled) on species yield was then computed. A variable is said to get a “hit” when its importance is greater than the greatest importance of the duplicated variables. This procedure was applied 100 times and a variable was selected if reaching a number of “hits” greater than the 95 % quantile of a binomial distribution Bin(100,0.5). Finally, given that i) ten imputed datasets were associated with each species and that ii) a model was fitted on each imputed version, we retained only the explanatory variables selected by all the ten models for each species.

### Model evaluation

The models’ goodness of fit was evaluated using the Root Mean Square Error (RMSE), wh<u>ich was calculat</u>ed for grain yield, for each model and imputed dataset as *RMSE* = 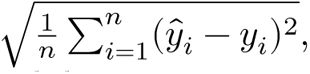, with *y*_*i*_ being the observed values and *y̑*_*i*_ the fitted values. The cross-validation error was assessed by splitting each dataset into four folds, ensuring that each fold includes at least one experimental unit from each experiment (as mixed-effect models cannot predict values on new modalities). First, the model’s performance was compared, using cross validation and without variable selection procedure. We then reported the performance of the MERF model, with the variable selection. The impact of the imputation process on overall variability was assessed by comparing the variance among imputations to the total variance of the residuals. Specifically, the proportion of variance due to the imputation process was computed as follows:

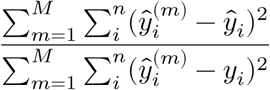

where *M* is the number of models that have converged over all the imputations, *y*_*i*_ is the *i*-th observation, *y̑_i_*^(*m*)^ is the fitted value for imputation *m*, 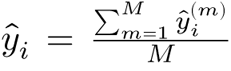 is the average fitted value between the *M* imputations (for details, see Mahmoud, 2023).

### Software

Data processing, statistical analysis, visualization, and reporting were performed with the R software.

We used R version 4.4.0 (R Core Team, 2024) and the following R packages: Boruta v. 8.0.0 (Kursa and Rudnicki, 2010a), ggh4x v. 0.2.8 (van den Brand, 2024), glue v. 1.7.0 (Hester and Bryan, 2024), here v. 1.0.1 (Müller, 2020), kableExtra v. 1.4.0 (Zhu, 2024), knitr v. 1.48 (Xie, 2014, 2015, 2024), latex2exp v. 0.9.6 (Meschiari, 2022), LongituRF v. 0.9 (Capitaine, 2020), nlme v. 3.1.164 (Pinheiro and Bates, 2000; Pinheiro et al., 2023), patchwork v. 1.2.0 (Pedersen, 2024), randomForest v. 4.7.1.1 (Liaw and Wiener, 2002), rmarkdown v. 2.28 (Xie et al., 2018, 2020; Allaire et al., 2024), rprojroot v. 2.0.4 (Müller, 2023), rsample v. 1.2.1 (Frick et al., 2024), tidyverse v. 2.0.0 (Wickham et al., 2019), viridis v. 0.6.5 (Garnier et al., 2024).

## Results

### Model evaluation

The multiple imputation procedure enabled for retaining 25 % of the experimental units in the dataset, preventing their removal, which would have occurred if rows with at least one missing value had been deleted. Moreover, imputing missing data did not significantly impact the fitted values. Specifically, the proportion of variance attributed to the imputation process ranged from 2.5 to 4.5 %. Consequently, the RMSE values reported here were averaged over the 10 imputed dataset.

The mean fitting abilities of the three types of models (LME, RF-based models without variable selection) were good and overall equivalent on the training dataset (RMSE = 0.2, 0.18, 0.16 t.ha^-1^, respectively for LME, RF and MERF). However, the two RF-based models performed much better than LME on the test dataset through the cross-validation procedure (RMSE = 0.44 t.ha^-1^ for both RF and MERF, ranging from 0.32-0.58) than the LME models (RMSE = 0.87 t.ha^-1^, ranging from 0.58-1.11). These results highlight the benefits of RF-based models, particularly in their handling of non-linear effects. Once accounting for variable selection with the Boruta method, we reported no differences between the RF and MERF models on the test dataset (RMSE = 0.44 t.ha^-1^).

**Figure 2.**
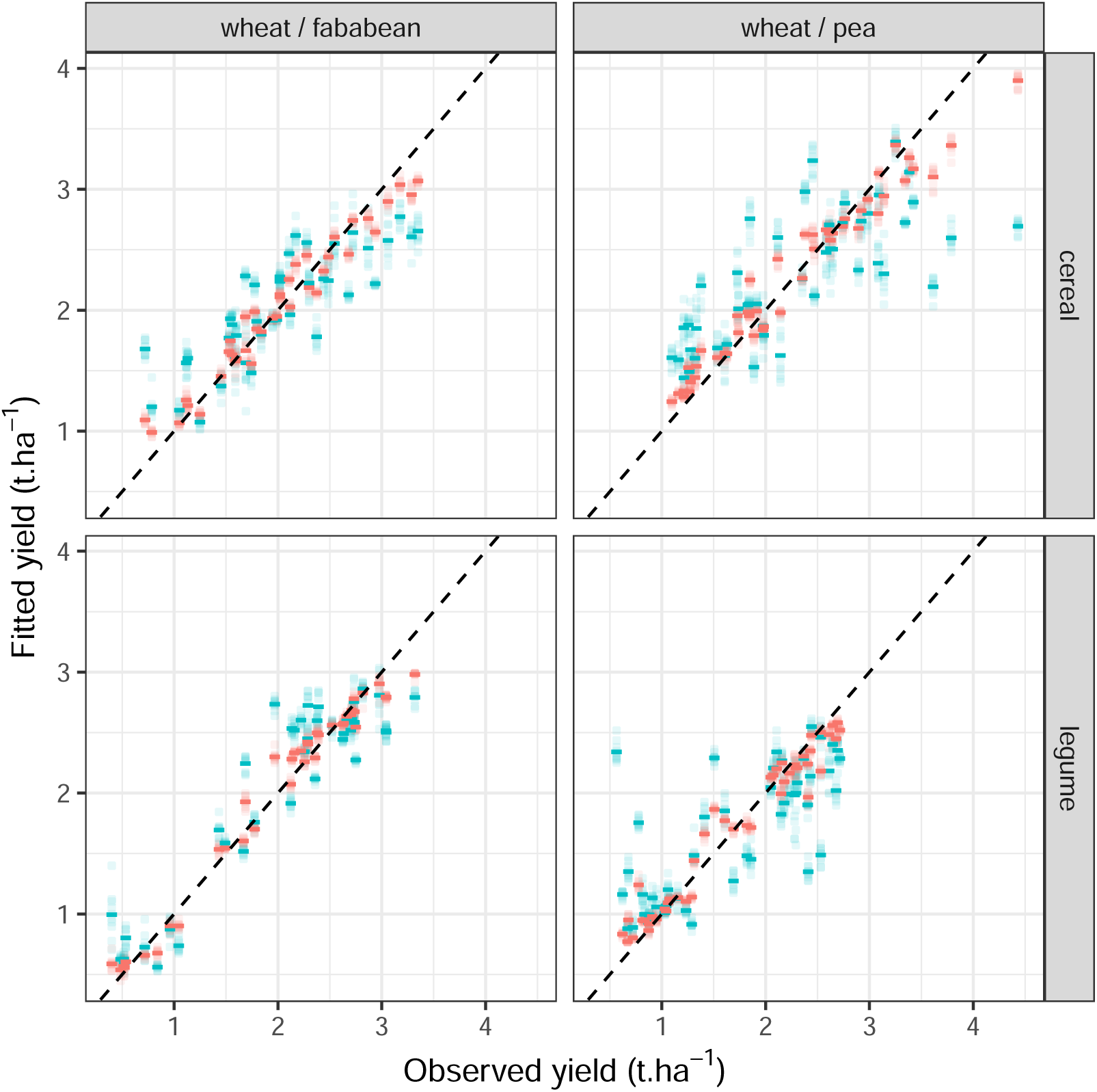
Fitted values as a function of observed values. The plot shows the relationship between fitted and observed values for different mixtures and their components. Each point represents an imputed dataset (10 in total), with the mean values shown as lines. The red lines indicate the model fitted using all available data, while the blue lines represent the model’s accuracy estimated using cross-validation. The mixtures are displayed in columns, and their components are arranged in rows.

The imputation procedure did not contribute significantly (1.8 % in average) to the variance of random effects in MERF models. For all species, the confidence intervals encompass 0, indicating no major effect of the experiment (Figure 3). Only in one case (faba bean in durum wheat / faba bean mixture), one experiment (2012) was associated with a negative impact on yield.

**Figure 3.**
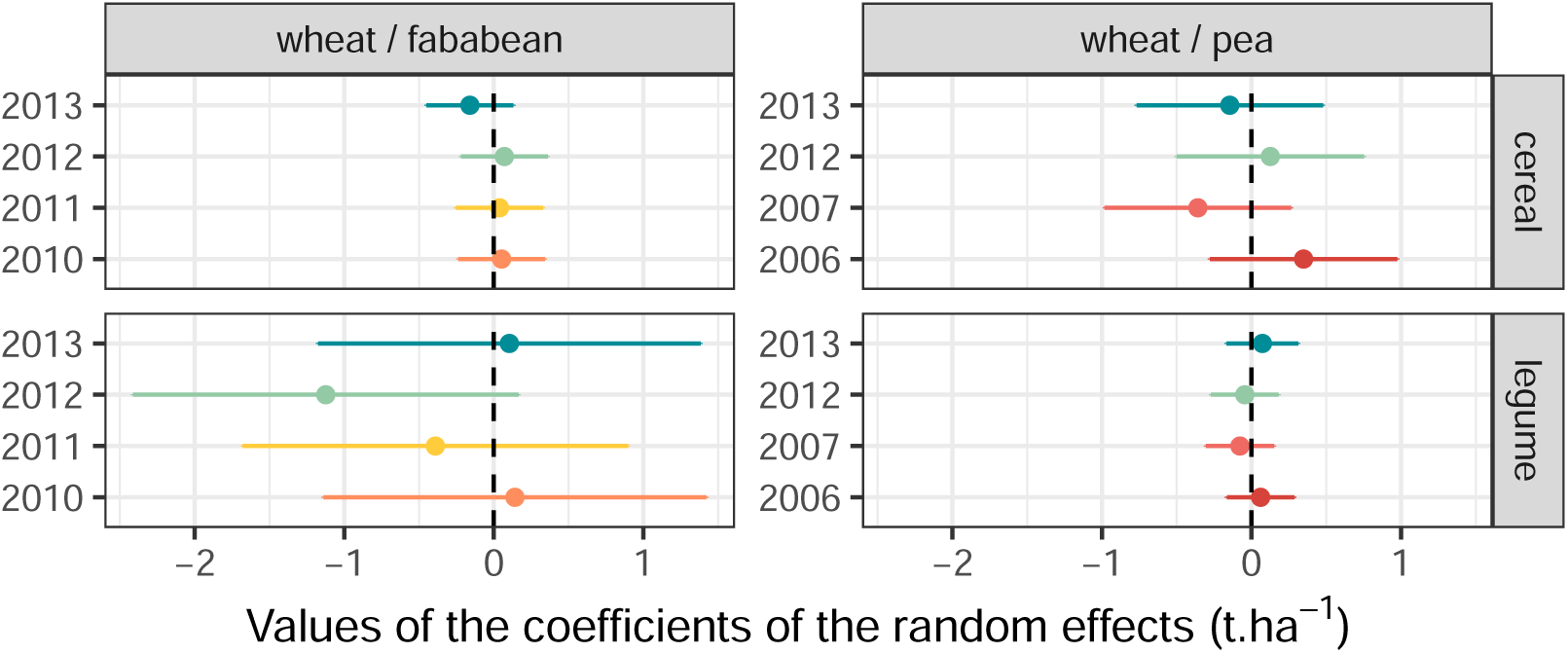
Mean values and confidence intervals of the coefficients of the random effect of MERF models. Error bars represent the confidence intervals 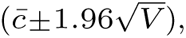, where *c̅* is the average random effect coefficient across imputations and is the variance of the random effect coefficient, in t.ha^-1^.

### Main plant-related features explaining species yield in mixture

We used a two-step approach to interpret our models (here, the MERF models) and identify the key features influencing intercropped species yield. First, we reduced the set of features by applying a stringent variable selection process during model fitting (Kursa and Rudnicki, 2010b). Then, we ranked the selected variables based on their contribution to the variance of the species yield (variable importance).

The variable selection process was both parsimonious and robust. Across species, the number of selected explanatory variables ranged from 4 to 7 (Figure 4), representing a significant reduction from the initial inclusion of 16 variables in each model. Specifically, we retained only the explanatory variables that were consistently selected across all 10 imputed datasets within each species model group.

**Figure 4.**
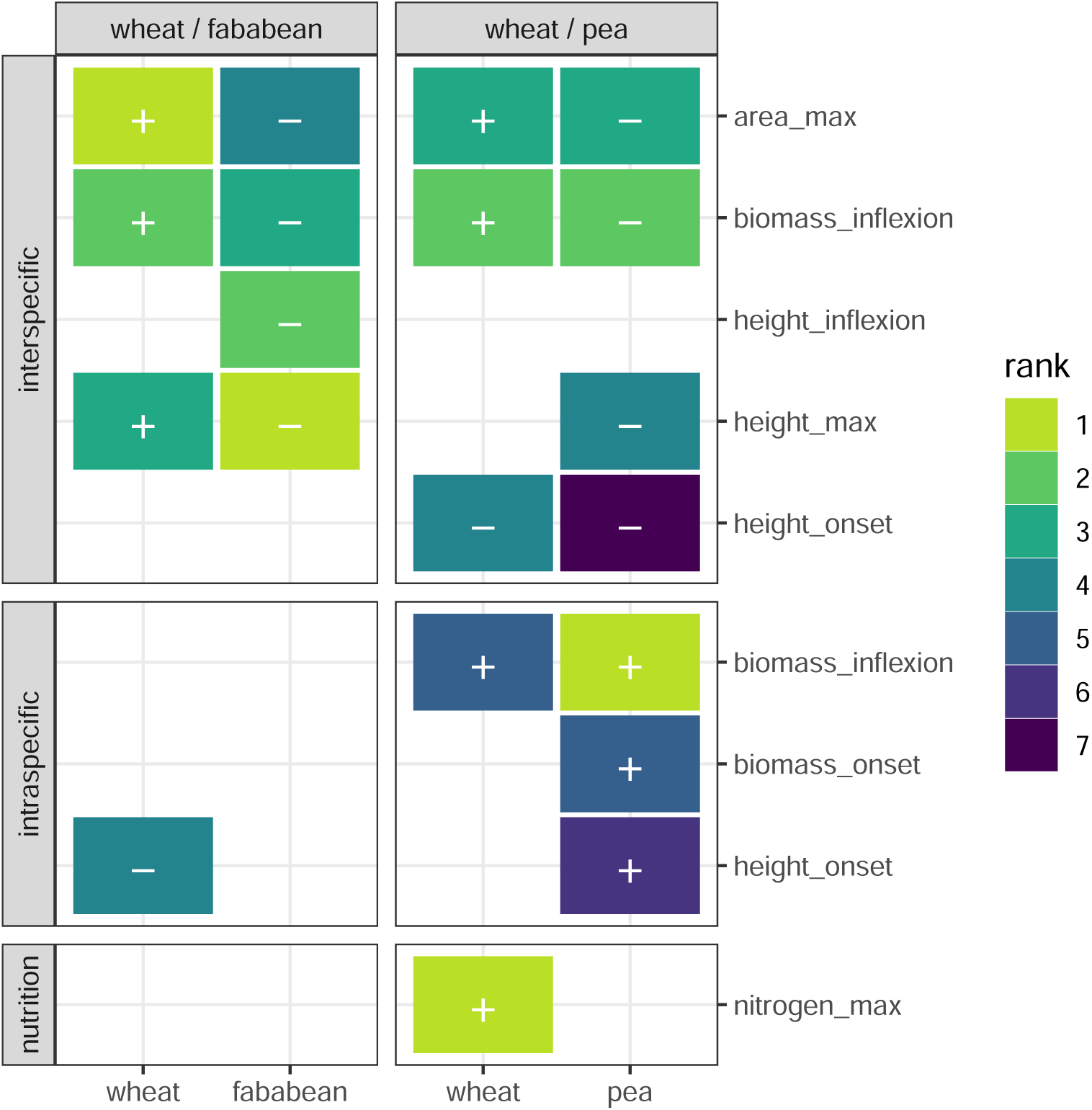
Variable importance in the two mixtures. The figure summarises the explanatory variables selected in the four MERF models, for each mixture component (in columns). The explanatory variables are organized in rows, and grouped into three types: i) interspecific indicators, calculated as the difference between the values of cereal and legume species; ii) intraspecific indicators, calculated as the difference between values in mixture and sole cropping conditions; and iii) agricultural practices, where only N status was selected. The colors indicate the ranking of the variables within the models, based on computed variable importance (with 1 being the most important). The sign (+/-) within each rectangle indicates a positive or negative Kendall correlation coefficient between the response variable (yield) and the explanatory variable.

Overall, interspecific differences within mixtures emerged as important across all mixture combinations (Figure 4). Among these, indicators related to biomass accumulation, maxi-mum LAI and height, which are proxies for species competitiveness, were particularly impor-tant in both mixtures. In contrast, intraspecific differences were generally less important, except for pea in wheat/pea intercrops.

The yield of a focal species in a mixture was positively correlated with its dominance, as reflected by its stature (height, leaf area, biomass), at the expense of the associated species (Figure 4 and 5). As a result, the same interspecific variables were selected for both wheat and the associated legume, but these variables had opposite effects on their respective yields. Specifically, wheat yield was highly and positively correlated with interspecific differences, while legume yield showed the opposite trend (Figure 5). These opposing relationships suggest that competition plays a crucial role in wheat/legume intercrops, with both species showing similar sensitivity to it. This is evidenced by the nearly identical absolute values of the correlation coefficients (mean *r* = 0.66, ranging from *r* = 0.23 to *r* = 0.83), with very close values for both species when considering a given feature and a relationship centered around zero for both species (Figure 5). when analyzed at the scale of the experiments, we observed overall lower correlation between performance and trait distance when factoring years (*r* = 0.53, ranging from 0.12 to 0.77). In some cases, such as the relation between wheat yield and interspecific maximum height distance, we observed sign-changing correlations between years (from *r* = −0.57 to *r* = 0.63), while the global relationship was positively correlated (*r* = 0.60).

**Figure 5.**
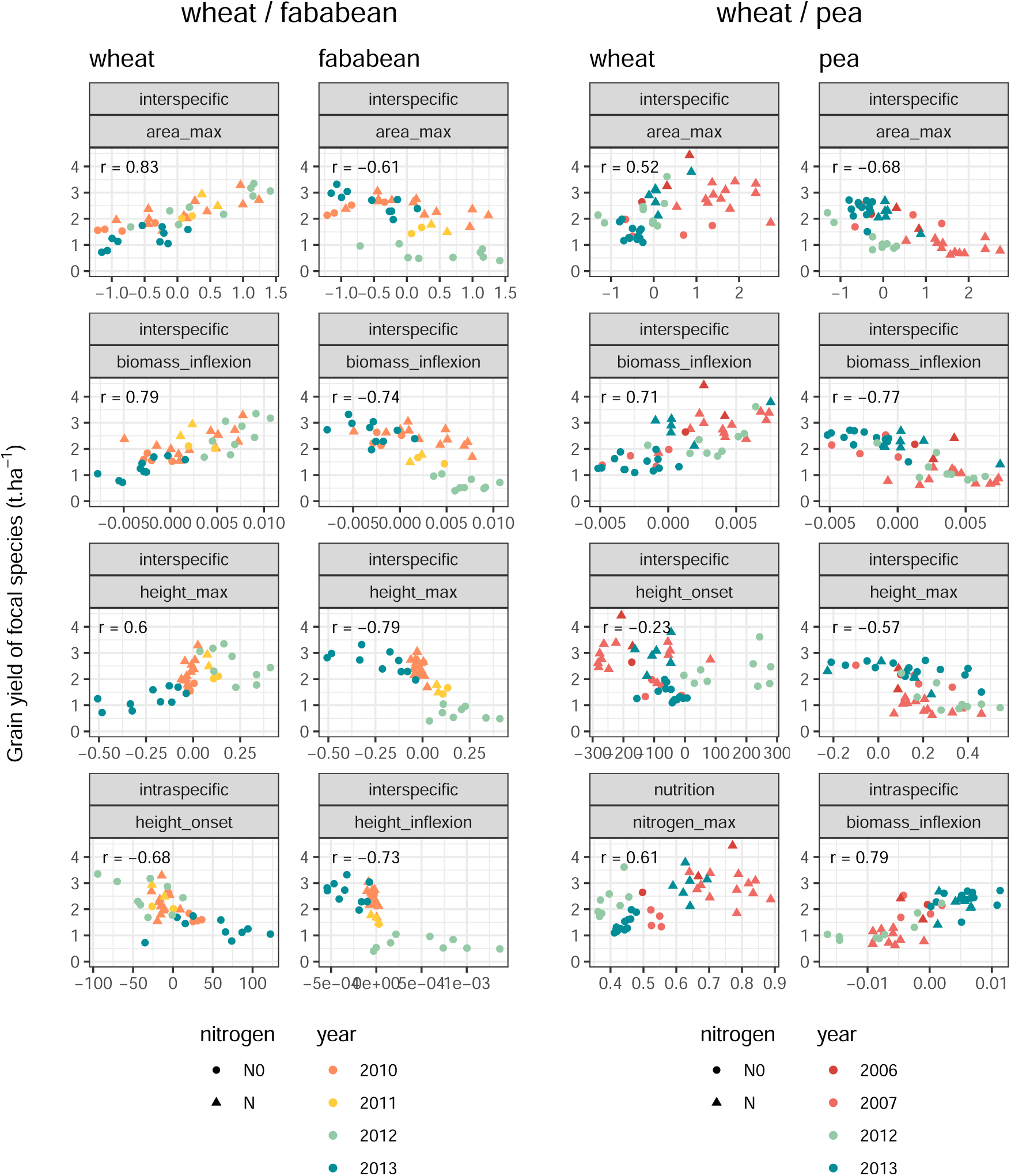
Bivariate relationships between grain yield and selected explanatory variables. Bivariate relationships between the selected explanatory variables (first four ranked variables), organized by mixture components (columns). Each point corresponds to an experimental unit (unique combination of {year, crop management}). The colour of the points encode the years (or experiments) and the shape encode the two N-fertilization levels, *i.e.* N0 = no fertilization, N = fertilization.

While the effects of competitive processes were consistent across all mixtures, some results were species-specific. In wheat/faba bean mixtures, wheat yield was partially explained by differences in the timing of its elongation phase (Figure 5, fourth row), suggesting a potential impact of intercropping on wheat phenology. Specifically, when wheat began its growth earlier in intercropping compared to sole-cropping conditions (a pattern observed in all experiments except in 2013), its yield was positively influenced. This earlier growth allows wheat to avoid some of the competition for resources with faba bean. A potential explanation for this could involve plant-plant signaling, mediated by light quality. Studies have shown that plants can perceived changes in light environment, prompting early developmental shifts, as seen in wheat/maize (Zhu et al., 2014), soybean/maize (Yang et al., 2014), and soft wheat/faba bean (Wang et al., 2024a) intercrops. In wheat/pea mixtures, several intraspecific indicators were identified for explaining pea yield, particularly the intraspecific difference in biomass growth rate between intercropping and sole-cropping conditions (Figure 4). It seems that experimental conditions (*i.e*, years) shaped and structured this relationship (Figure 5, col. 4, row 4). Specifically, when the pea was able to grow faster in intercropping than in sole-cropping, its yield was positively affected.

### Impact of agricultural-relaed features importance in mixtures

Fertilization, as indicated by crop N status (NNI), was identified as the main explanatory variable in only one case: for wheat in wheat/pea intercrops (Figure 4). In this context, wheat yield was positively influenced by its N status, with a notable yield gain resulting from a moderate N amount (Figure 5, col. 3, row 4). A threshold effect was observed, with yield gain occurring at NNI values above 0.6, which corresponded to the N-fertilized experimental units. Consistent with previous studies, N-fertilization affected the balance between the two intercropped species (Pelzer et al., 2012; Duchene et al., 2017; Mahmoud et al., 2022). This was further evidenced by changes in the interspecific differences in maximum LAI and shoot biomass growth rate across varying levels of N-fertilization (Figure 5, col. 3, rows 1 and 2).

In all models, cultivar identity was never selected as a key explanatory variable, likely because it was indirectly accounted for by variables representing plant-plant interactions. However, it is challenging to separate the effects of cultivar identity from other experimental factors due to the study’s experimental design (Table 1). When focusing on the cultivars used, the years 2010 and 2011 were quite similar for wheat/faba bean mixtures, as were 2006 and 2007 for wheat/pea mixtures. Only 2012 and 2013 included a greater diversity of cultivars for all the three species.

In the relationships between species yield and variables related to biomass or leaf area, the statistical individuals were not clustered according to year or N-fertilization factors (Figure 5). However, height-related variables (Figure 5, row 3) revealed a clearer clustering by experiments/year, particularly for wheat/faba bean mixtures. This clustering cannot be attributed to cultivar effects, as the same cultivars were used in both 2012 and 2013. Instead, it is likely driven by significant differences in precipitation, with 2013 experiencing around 80% more rainfall between crop sowing and harvest, compared to other years (Table 1). While notable trends between yield and height-related variables emerged when combining several experiments, these trends were not consistently observed within individual experiments.

## Discussion

### Merging statistical cultures to balance predictive accuracy and model complexity

#### Performance of linear vs RF-based models

As expected, the performance gap between linear and random forest (RF)-based models indicated that the relationships between species yield and explanatory variables were better captured by machine learning algorithms (Breiman, 2001b). However, the goal of accounting for the dependence of observations within the dataset was only partially achieved. The confidence intervals around the random effect coefficients suggested that not all variability associated with the experimental factor was captured in the random effect. This was highlighted by the similar performance observed in the RF and MERF models, as well as the grouping structure observed in the bivariate relationships between species yield and explanatory variables.

#### Taking into account environmental effects

This difficulty of random-effects models in capturing all intergroup variability has already been pointed out (Gelman and Hill, 2006). To better estimate the random effects coefficients, it would be interesting to include a heteroscedastic structure in the mixed model. This could more accurately account for within-experiment dependencies but would require additional experimental units per experiment to estimate the variance correctly (Bolker et al., 2009). In previous iterations of this work (Mahmoud, 2023), accounting for the environment through physical variables led to weaker predictive approaches compared to the one presented in the current study, whether it was done by: i) classifying experiments into homogeneous climate groups (envirotyping, Chenu et al., 2013), or ii) identifying key moments during the growth cycle when climatic variables influenced yield (Picheny et al., 2019).

We assume that the use of plant traits as predictors already captures a significant portion of the environmental effects. Considering plant traits (and trait-derived variables) as a signature of the processes driving mixture functioning provides a robust basis for predictive models in agricultural conditions. However, increasing experimental variability -for example, by including more diverse geographic locations -could challenge this assumption. In our design, experiments were conducted in contrasting years but at the same location. We argue that greater geographic variability would likely widen the performance gap in favor of MERF models.

#### Integration of mechanistic and machine learning models

Machine learning models, including RF, have shown remarkable success across scientific disciplines, extending beyond statistics to fields such as ecology and social sciences (Jordan and Mitchell, 2015). Over time, the line between the different modeling approaches has blurred, leading to more hybrid methods (Zhao, 2021). Each type of modeling — mechanistic, data-driven, and algorithmic — comes with its own trade-offs, such as flexibility, causality, predictive accuracy, and simplicity. Researchers should position their modeling approach within this broader *space of models* (Miller et al., 2021).

This is particularly relevant in agroecology, where agrosystems are becoming increasingly complex, while models are still mainly developed per species or per type of modeling school (Gaudio et al., 2022). We argue for more hybrid modeling approaches, aiming to describe systems through a combination of models, rather than using a single model with a inevitably limited domain of validity.

#### Challenges of random forest in variable selection and dataset imbalance

While RFs have strong predictive ability and robustness, they do not inherently account for the clustered data structure. We used the RF algorithm in MERF not with the goal of prediction, but to identify relevant predictors from a set of candidate variables. In this context, the developed models proved to be suitable for the intended purpose of using proxies to capture the plant interactions occurring within the studied mixtures. While our approach appears to be scalable with more experiments and predictors, two limitations arise. First, using RF as a variable selection tool becomes more challenging as the number of predictors increases. Second, as datasets become larger, importance-based methods may struggle with the increased imbalance of experimental designs.

When using RF mainly to select variables rather than predict, Strobl et al. (2007) mentioned that the value and interpretation of the variable importance measure must actually depict the importance of the variable and should not be affected by any other characteristics. They showed that the importance estimations in the RF algorithm were not reliable if the set of predictors included categorical variables with large variations in their category number along with continuous variables. In our use case, our predictor set was not falling under those conditions: they were mostly continuous, with only one categorical variable with a low number of categories (the cultivar identity, with 5 modalities maximum).

Regarding dataset imbalance, larger datasets may introduce more confounding effects (Mahmoud et al., 2024), warranting future research into causal relationships (Pearl, 2009), especially between traits. For example, light interception as a driver of biomass accumulation is a relevant causal relationship. Causal relationships are of great significance in ecology (Arif and Macneil, 2022), and should be further explored in intercropping studies.

### Predictive approaches for crop ecology

#### Interspecific differences play a predominant role in explaining intercropped species yield

Our modeling approach effectively explained species yield in mixtures using only 4-7 explanatory variables. Compared to mechanistic models focused on intercrops (Gaudio et al., 2019), mixed-effects random forest models were more parsimonious while maintaining strong explanatory power. The process of selecting the most influential variables could also benefit the design of mechanistic models (e.g. Casadebaig et al., 2016, for wheat in sole crop), as it helps to prioritize the key variables and processes that are most important in explaining mixture functioning.

In all fitted models, features indicative of interspecific plant-plant interactions were selected more frequently than those comparing mixtures to sole cropping. Particularly, differences in growth dynamics and final stature between species were crucial in explaining their respective yields, confirming the role of plant interactions within crop mixtures (Justes et al., 2021). The description of growth processes (onset, rate, maximum) proved particularly useful, highlighting the importance of capturing dynamic measurements in agronomic experiments, taking into account variables routinely measured in field trials (biomass, height, leaf area).

Dialogue between modelers and experimenters is crucial, fostering a two-way exchange that refines both model predictions and experimental designs (Craufurd et al., 2013; Rötter et al., 2018). This iterative feedback loop helps guide experiments towards collecting the most relevant data, improving the models’ predictive capacity, and enabling more targeted field measurements. Thus, this study provides valuable feedback to experimenters, upon whom the development of accurate, data-driven models was highly dependent.

The difference in shoot biomass growth rates between intercropped species was identified as a key factor in all four models, consistent with findings from other studies on different intercrops such as oat / lupin, rapeseed / maize, or rapeseed / soybean (Dong et al., 2018; Engbersen et al., 2021). Interestingly, while several studies highlighted the importance of differences in growth biomass onset to explain intercropped species yield (Dong et al., 2018), this feature was not selected in our models. However, this discrepancy may be explained by the fact that many studies emphasizing biomass onset deal with relay intercrops, where species are sown at different times, relying on temporal niche complementarity (Yu et al., 2015). Our study focused on simultaneous sowing, where such temporal dynamics may be less relevant.

In contrast to interspecific plant-plant interactions, we found that it is unlikely to explain the functioning of the mixtures based on species behavior in the sole crop, as few intraspecific variables were selected. This observation supports the need for breeding programs specifically tailored to mixed cropping systems, rather than relying on traits selected in sole crops (Moore et al., 2022). However, we did not include trait differences between the legume and the cereal grown in sole crop in our models, which may be worth exploring further as it may provide additional insight into crop mixture performance.

#### Relationships between performance and trait distance are symmetric for both intercropped species

According to the literature, relationships between yield and the relative trait distance were expected (Kunstler et al., 2012), without assumptions about their shape and strength. Our results highlighted that the relationships between a species’ yield and relative biomass distance were globally linear and inversely related for the two intercropped species, indicating that the dominance of one species occurred at the expense of the other (Mahaut et al., 2023).

When a variable was selected for both species in the mixture, the strength of the correlations was similar for both. This symmetry was unexpected. Cereals are typically considered more competitive than legumes as they tend to accumulate biomass faster and use resources more efficiently (Hauggaard-Nielsen et al., 2008; Lithourgidis et al., 2011), which should theoretically give them an advantage in mixture conditions. Our results indicate that it’s unlikely that a single mixture can optimize yields for both species simultaneously. In practical terms, this means farmers and researchers need to consider trade-offs when designing intercropping systems.

Depending on the farmer’s objectives when cultivating intercrops, the goal may be to promote the dominance of a particular species (*e.g.*, for the legume Podgorska-Lesiak and Sobkowicz, 2013; Viguier et al., 2018) or to aim for a balance between the two species (Hauggaard-Nielsen et al., 2008; Bedoussac et al., 2015). This study shows that, with the same pair of species, all these outcomes are equally possible. To move towards application, we thus think future research should focus on how a state of equilibrium or dominance occurs as a function of species or cultivar combinations, given cropping conditions. To limit the number of combinations of considered mixtures, cultivars could be clustered into functional groups (Montazeaud et al., 2020) with a modeling step at this grain.

#### Relationships between performance and trait distance are scale-dependent

In this study, we identified significant bivariate relationships between yield and explanatory variables when aggregating all experiments. These findings suggest overarching trends that emerge when a wide range of environmental conditions and management practices are considered together. However, these trends were not always consistent when down-scaling at the individual experiment level, suggesting that local factors (such as soil properties, climate conditions, or agricultural management) can override or mask broader trends at smaller scales, as already illustrated in other scientific fields (*e.g.*, Scheiner et al., 2000 in ecology, on the relationship between species richness and productivity).

This discrepancy between global and local results highlights the complexity of agronomic systems, where covariation between variables may not behave uniformly across all contexts. The variability observed at the local level could be driven by site-specific factors or other unmeasured interactions that influence crop responses in ways not fully captured by linear predictors alone. Therefore, while aggregated analyses provide valuable insights into generalizable trends, they should be applied with caution in local contexts (Scheiner et al., 2000), *i.e.*, covariation at one scale cannot necessarily be interpreted as indicative of a process at another scale (Pollet et al., 2014). Thus, in our case, the linear relationships between species yield and fitness differences (Wang et al., 2024b) were evident. The challenging issue lies in identifying which local factors determined the positioning of a species yield along this relationship.

## Conclusion

Crop science can greatly benefit from aggregating distinct experimental datasets into global datasets (Mahmoud et al., 2024). This approach decreases the cost of data (reuse), increases the reproducibility of studies (Lowndes et al., 2017), and enables the testing of findings across multiple experiments through joint analysis. In this study, we proposed an original modeling approach that balances these goals: using machine learning to select and assess how species-level traits contribute to community productivity, while emphasizing on accounting for ecological knowledge (with tailored predictors) and environmental variance (using mixed-effects). Ultimately, as pointed out by Enquist et al. (2024), such model-centered approaches could gradually evolve both towards generalisation or specialisation, reflecting a key tension in generating knowledge about agrosystems. On one hand, models could be refined to incorporate details highlighted by theoretical ecology, increasing their applicability domain.

On the other hand, new open datasets could be curated to evaluate model performance, providing feedback by identifying specific contexts or outliers where existing theories fail.

## Author Contributions

analysis: PC, NG, XG, NH, RM writing: PC, NG, XG, NH, RM

coordination: PC, NG, NH

funding: PC, NG

all authors commented on previous versions of the manuscript

all authors read and approved the final manuscript

## Data Availability

This repository contains the processed data, the model fitting procedure, and the code to produce the manuscript and the figures.

